# Targeting RNA Polymerase I transcription synergises with TOP1 inhibition in potentiating the DNA damage response in high-grade serous ovarian cancer

**DOI:** 10.1101/849307

**Authors:** Shunfei Yan, Piyush B. Madhamshettiwar, Kaylene J. Simpson, Sarah Ellis, Jian Kang, Carleen Cullinane, Karen E. Sheppard, Katherine M. Hannan, Ross D. Hannan, Elaine Sanij, Richard B. Pearson, Keefe T. Chan

## Abstract

Limited effective therapeutic options are available for patients with recurrent high-grade serous carcinoma (HGSC), the most common histological subtype accounting for the majority of ovarian cancer deaths. We have shown efficacy in poly-ADP ribose polymerase (PARP) inhibitor-resistant HGSC for the RNA Polymerase I (Pol I) transcription inhibitor CX-5461 through its ability to activate a nucleolar-associated DNA damage response (DDR). Here, we screen the protein-coding genome to identify potential targets whose inhibition enhances the efficacy of CX-5461. We identify a network of cooperating inhibitory interactions, including components of homologous recombination (HR) DNA repair and DNA topoisomerase 1 (TOP1). We highlight that CX-5461 combined with topotecan, a TOP1 inhibitor used as salvage therapy in HGSC, induces robust cell cycle arrest and cell death in a panel of HR-proficient HGSC cell lines. The combination potentiates a nucleolar-associated DDR via recruitment of phosphorylated replication protein A (RPA) and ataxia telangiectasia and Rad3 related protein (ATR). CX-5461 plus low-dose topotecan cooperate to potently inhibit xenograft tumour growth, indicating the potential for this strategy to improve salvage therapeutic regimens to treat HGSC.

## Introduction

High-grade serous carcinoma (HGSC) is the most prevalent histological subtype of epithelial ovarian cancer with the worst prognosis. HGSC is characterised by nearly universal *TP53* mutations (>96%), and up to 50% of HGSC harbour defects in homologous recombination (HR) DNA repair genes, including breast cancer related antigen (*BRCA)1* and *BRCA2*, which are necessary for maintaining genomic integrity^1^. Defects in DNA repair confer exquisite sensitivity of HGSC to standard DNA-damaging chemotherapies (carboplatin/cisplatin and paclitaxel) as well as the recently FDA-approved poly-ADP ribose polymerase (PARP) inhibitors (olaparib, rucaparib, niraparib, talazoparib), which are synthetic lethal by generating catastrophic DNA damage and cell death in cells lacking HR ^2^. However, resistance to these therapies frequently develops via multiple mechanisms, including restoration of HR leading to disease relapse ^3^, highlighting the need to identify new therapeutic targets to overcome resistance. HR-proficient HGSCs including the 20% harbouring *CCNE1* amplifications are associated with intrinsic platinum-based chemotherapy resistance and poor clinical outcome ^4^. Despite few available salvage therapies, the DNA topoisomerase 1 (TOP1) inhibitor topotecan has shown promise, particularly when combined with the angiogenesis inhibitor Sorafenib in platinum-resistant ovarian cancer ^5^.

Increased activation of key oncogenic signalling pathways (PI3K/AKT, RAS/MAPK and MYC) upstream of ribosome biogenesis constitutes an additional hallmark of HGSC ^6^, and we hypothesise that inhibiting ribosome biogenesis with the ribosomal RNA gene (rDNA) transcription inhibitor CX-5461 can provide an effective cancer therapeutic option ^7^. Indeed, we have shown encouraging responses in a Phase I clinical trial in haematological malignancies ^8^, and Canadian trials in breast cancer are ongoing ^9^. We have demonstrated in cells with intact p53, CX-5461 induces an impaired ribosome biogenesis checkpoint leading to apoptosis, cell cycle arrest or senescence; however, in cells with inactivated p53 such as HGSC, CX-5461 induces the DNA damage response (DDR) pathway ^10^, which is emerging as an attractive clinical target due to the high levels of genomic instability characteristic of HGSC ^11^.

We have demonstrated CX-5461 activates the DDR at the nucleoli, the sites of RNA Polymerase I (Pol I) transcription, and exhibits therapeutic efficacy in both PARP inhibitor-sensitive and -resistant HGSC ^12^. To uncover additional combination therapies exploiting the non-canonical nucleolar-associated DDR and identify potential targets whose inhibition can enhance the efficacy of CX-5461, we performed a whole protein-coding genome RNAi screen in p53-mutant OVCAR4 cells, which closely resemble the genomic profile of HGSC ^13^. We confirmed HR deficiency sensitises HGSC to CX-5461. We also identified that knockdown of TOP1 or treatment with the TOP1 inhibitor topotecan synergises with CX-5461 in multiple HR-proficient HGSC cell lines. Topotecan treatment in the clinic is limited by significant toxicity including myelosuppression ^14^; however, when used in low doses with CX-5461, the combination induces cell death and cell cycle arrest by potentiating the nucleolar DDR without inducing DNA resection. We identify a new therapeutic approach to target HGSC while minimizing the toxicities normally associated with standard of care DNA-damaging therapies.

## Materials and Methods

### Cell culture

Human HGSC cell lines (OVCAR3, OVCAR4 and CAOV3) were obtained from the National Cancer Institute. All cell lines were short tandem repeat (STR) characterised against American Tissue Type Collection (ATCC) or ExPASy databases to ensure authenticity of origin. Mycoplasma tests were performed routinely by PCR. All cells were cultured in RPMI-1640 media (Gibco) supplemented with 10% Fetal Bovine Serum (FBS) (Sigma-Aldrich) and 2mM GlutaMax™ (Gibco) at 37°C and 5% CO_2_.

### Reagents and antibodies

CX-5461 was provided by SYNkinase and prepared in 50 mM NaH_2_PO_4_. Topotecan (Hycamtin^®^, Novartis) was obtained from the Peter Mac pharmacy and was dissolved in 0.9% saline. DBL^™^ doxorubicin hydrochloride injection was purchased from Hospira and diluted in phosphate buffered saline (PBS). The pan-caspase inhibitor Q-VD-OPh (Cat #1901) was purchased from APExBIO and dissolved in dimethyl sulfoxide (DMSO). A list of antibodies used in this study is provided in Supplementary Table S1.

### Genome-wide protein-coding RNAi primary screen and analysis

On a screening day, 160 μL DharmaFECT 4 (Horizon Discovery) was mixed with 50 mL Opti-MEM^®^ (Gibco) (sufficient for 16 assay plates), and 44 μL lipid:Opti-MEM mixture was then aliquotted to each well of a 384-well black-walled plate (Corning, Cat. #: 3712) containing 6 μL 1 μM SMARTpool siRNA (Horizon Discovery) using a BioTek EL406™ washer dispenser (final SMARTpool concentration 40 nM). The transfection mixture was mixed and complexed for 20 minutes, and 12.5 μL mixture was then aliquotted into three plates (4 replicate plates in total) using a Caliper Sciclone ALH3000 liquid handler. During this period, OVCAR4 cells were trypsinised and resuspended at 5.6 × 10^4^ cells/mL. Twenty-five μL OVCAR4 cells (1400 cells) were then dispensed into 384-well plates with the final concentration of the SMARTpool siRNA at 40 nM, and plates were incubated at 37°C and 5% CO_2_. At 24 h post-transfection, the medium was replaced with cell culture medium containing either 80 nM CX-5461 or 400 nM NaH_2_PO_4_. Cells were incubated for another 48 h and fixed with 2% PFA in PBS for 10 min, followed by permeabilisation with 0.3% TritonX-100 in PBS for 10 min. Cells were washed with PBS, stained with 100 ng/mL DAPI for 20 min. Cells were washed twice with PBS and 25 fields were imaged using the ArrayScan VTI high-content system (Thermo Fisher Scientific) with a 20x/0.4 NA objective and an ORCA-ER camera and a 5-ms exposure time. The Cellomics Morphology V4 Bioapplication was used to analyse cell number as determined by DAPI staining.

To identify synergistic gene candidates, we used a combination of the difference in relative cell number between vehicle- and CX-5461-treated target siRNA plates normalised to ON-TARGETplus (Horizon Discovery) non-targeting control (siControl) ≥ 0.25, and a value for Bliss independence ≤ 0.9, which was calculated as follows ^15^.

Bliss Independence = (Relative cell number (siRNA + CX-5461)) / (Relative cell number (siRNA + vehicle) × Relative cell number (siControl + CX-5461))

### Secondary deconvolution screen

Based on the above criteria, we selected 372 genes for a secondary deconvolution screen using the 4 individual siRNA duplexes that comprised the SMARTpool arrayed separately (1 duplex/well). For the secondary screen, the 1 μM SMARTpool siRNAs were replaced by 0.45 μM individual siRNA duplexes for a final concentration of 25 nM/duplex. We modified our selection criteria to define high-confidence hits as those with a difference in relative cell number between vehicle- and CX-5461-treated target siRNA plates normalised to siControl ≥ 0.15 and Bliss Independence ≤ 0.8. We classified genes with ≥ 2 high-confidence hits as those displaying synergy.

### qPCR

The RNeasy^®^ Mini Kit (Qiagen) was used to extract total RNA as per the manufacturer’s instructions. One hundred ng of total RNA was used as a template for cDNA synthesis using SuperScript^™^ III reverse transcriptase (Invitrogen^™^ #18080093), hexameric random primers and dNTPs. Quantitative real-time PCR (qPCR) reactions were performed using Fast SYBR^®^ Green reagents in a StepOnePlus^™^ Real-Time PCR system (Applied Biosystems^™^) with a +0.7^°^C melt increment. RNase-free water was used as a negative control. Changes in target gene expression were normalised to *NONO* housekeeping gene and fold change was determined by using 2^(-ΔΔ*C_t_*). Primer sequences are listed in Supplementary Table S2.

### Immunoblotting

Cells were washed twice with PBS and whole cell lysates were prepared in Western solubilisation buffer (20 mM HEPES pH 7.9, 2% SDS, 0.5 mM EDTA). Protein was transferred to polyvinylidene fluoride (PVDF) membranes, which were blocked in 5% skim-milk TBS 0.1% Tween® 20 (TBST) for 45 min at RT. Membranes were incubated with primary antibodies overnight at 4°C, washed three times in TBST for 10 min, incubated with HRP-conjugated secondary antibodies for 1 h at room temperature and then washed. Membranes were visualised using Western Lightning^™^ Plus enhanced chemiluminescence (Perkin Elmer) by exposure to film (Fujifilm SuperRX) or imaged by a ChemiDoc^™^ Touch Imaging System (Bio-Rad Technology). Digital scans of film were acquired using an Epson Perfection V700 Photo at ≥ 300dpi.

### Immunofluorescence

Five thousand cells were seeded into 8-well Nunc™ Lab-Tek™ II Chamber Slides™ (Cat. #: 154534) per well. Cells were cultured for 72 h, followed by drug treatments at the indicated timepoints and doses together with 10 μM 5-ethynyl-2’-deoxyuridine (EdU) (Sigma-Aldrich, Cat. #900584). Cells were fixed in 4% paraformaldehyde and permeabilised with 0.5% Triton X-100 in PBS. EdU was fluorescently labelled with 0.5 μM Click-iT^™^ EdU Alexa Fluor^™^ 647 Azide (Invitrogen^™^ #A10277) in labelling buffer (100 mM Tris, pH 8.5, 100 mM ascorbic acid, 1 mM CuSO_4_) for 30 min at room temperature and subsequently stained with the indicated antibodies. Cell nuclei were stained with 500 ng/mL 4’,6-diamidino-2-phenylindole (DAPI) diluted in PBS. Slides were mounted with Vectashield^®^ Antifade Mounting Media (Vector Laboratories, Cat. #: H-1000). Images were captured by a Zeiss LSM 780 confocal microscope using a 20x/0.8 NA Plan-apochromat objective and were analysed with FIJI ^16^ and CellProfiler ^17^.

### Cell cycle analysis

Cells were labelled with 10 μM 5-bromo-2′-deoxyuridine (BrdU) (Thermo Fisher Scientific, Cat. #: B23151) for 30 min, collected with supernatant and fixed with ice-cold 80% ethanol. Fixed cells were centrifuged, resuspended in 2N HCl + 0.5% Triton X-100 (v/v) and acid was neutralised with 0.1 M Na_2_B_4_O_7_·10H_2_O (pH 8.5). Cells (2 × 10^5^) were incubated with 100 μL anti-BrdU antibody (0.5 μg/mL) in dilution buffer (PBS + 2% FBS) + 0.5% Tween-20 for 30 min at room temperature, washed with dilution buffer and resuspended in 100 μL in Alexa Fluor 488 donkey anti-mouse IgG (5 μg/mL) in dilution buffer + 0.5% Tween-20 for 30 min on ice, then washed. Cells were resuspended in 10 μg/mL propidium iodide (PI) (Sigma-Aldrich, Cat. #: P4170) in dilution buffer and analysed by flow cytometry on a BD FACSCanto™ II. Quantitation of cell cycle populations was performed using FlowJo software.

### Clonogenic assay

Cells (1 × 10^5^/well) were seeded into 6-well plates (BD Falcon). Cells were cultured for 24 h, then drugged as indicated. At 48 h after drug treatment, the drugs were removed by washing twice with PBS. Cells were then cultured in normal medium for another 5 d when the vehicle-treated cells reached confluency. Cells were then fixed with 100% methanol for 10 min, stained with 0.1% crystal violet solution for 20 min, thoroughly washed with PBS and the stained plates were dried, imaged on an IncuCyte^®^ ZOOM (Essen Bioscience) using a 10x/0.3 NA objective and analysed for cell confluence.

### DNA comet assay

DNA comet assays were performed using the CometAssay^®^ Reagent Kit (Trevigen, Cat. #: 4250-050-K) according to the manufacturer’s protocol. Briefly, cells treated with indicated drugs were trypsinised and washed once with ice-cold PBS, and then resuspended in ice-cold PBS at 1 × 10^5^ cells/mL. Cells were mixed with molten low-melt agarose at 37°C at a ratio of 1:10 and 50 μL of the mixture was immediately pipetted onto a CometSlide™. The slides were incubated in the dark at 4°C for 30 min to solidify the agarose and immersed in 4°C Lysis Solution overnight. Slides were then immersed in freshly prepared Alkaline Unwinding Solution for 1 h at 4°C in the dark, and electrophoresed in 4°C Alkaline Electrophoresis Solution at 300 mA for 40 min. The slides were washed with ddH_2_O twice, followed by 70% ethanol for 5 min. Slides were dried at 37°C for 15 min, and then stained with 2.5 μg/mL PI in PBS for 30 min at room temperature. Slides were rinsed twice in ddH_2_O and completely dried at 37°C. Images were captured using a VS120 Virtual Slide Microscope (Olympus) using a 10x objective and analysed with the OpenComet ^18^ plugin for ImageJ.

### Xenograft transplantation

All animal studies were conducted according to the protocols approved by the Animal Experimentation Ethics Committee at the Peter MacCallum Cancer Centre. Non-obese diabetic severe-combined immunodeficiency gamma (NSG) mice (6-8 week-old female) were purchased from the Garvan Institute (Australian BioResources) and housed in animal cages under standard laboratory conditions. OVCAR3 cells (6 × 10^6^) in 100 μL ice-cold PBS:Matrigel (1:1) were subcutaneously injected into the right flank of mice anaesthetised with isoflurane. Tumour-bearing mice were weighed and measured twice weekly with callipers. Once tumours reached an average volume of 100 mm^3^, mice were randomised into 4 groups of 10 mice. Mice were dosed with vehicle (25 mM NaH_2_PO_4_) or CX-5461 (30 mg/kg) via oral gavage twice weekly while topotecan (5 mg/kg (experiment #1) or 2.5 mg/kg (experiment #2)) or vehicle (0.9% saline), were delivered via intraperitoneal injection twice weekly. Mice were dosed for 4 consecutive weeks for a total of 8 doses. Body weight and tumour volumes were monitored daily during and after the treatment period. Mice were euthanised by cervical dislocation upon reaching any ethical endpoint of the experiment (signs of distress or tumour volume ≥ 1200 mm^3^).

### Statistical analysis software

Prism 8 (GraphPad) was used for all the statistical analyses as indicated, including dose response curves, *t* tests, Wilcoxon rank-sum non-parametric tests, ordinary one-way ANOVA, Kruskal-Wallis nonparametric one-way ANOVA and Mantel-Cox tests. Synergy was quantified using Combenefit ^19^ (software based on the Bliss independence).

## Results

### A functional genomics screen identifies a network of genes that when depleted cooperates with CX-5461 to inhibit HGSC cell proliferation

The Pol I transcription inhibitor CX-5461 has efficacy in a PARP inhibitor-resistant patient-derived xenograft (PDX) model of HGSC ^12^. To further identify genes that, when depleted could synergise with CX-5461 to inhibit cell proliferation, we used human OVCAR4 HR-proficient HGSC cells in a protein-coding genome-wide RNAi screen (Fig. 1A). Based on the reduction in cell number relative to siControl and Bliss Independence to assess combinatorial effects, we identified 372 genes whose knockdown demonstrated synergy with CX-5461 (Table S3). Gene ontology analysis identified enrichment in several functional processes, with DNA damage:double-strand break (DSB) repair being the most significant (Fig. 1B, Table S4). Deconvolution screening using individual siRNA duplexes and more stringent metrics validated 20 genes using a concordance criterion of 2 out of 4 (Table S5). STRING network analysis of these genes demonstrated significance in homologous recombination (HR) (Fig. 1C), consistent with a previous study in breast cancer ^20^ and our findings that CX-5461 is synthetic lethal with HR deficiency and a HR deficiency gene expression signature predicts HGSC sensitivity to CX-5461 ^12^. Notably, treatment of OVCAR4 cells depleted of BRCA2 (4/4 siRNA duplexes validated) with CX-5461 led to significant growth inhibition (Fig. 1D), which was associated with micronuclei formation, suggesting enhanced genomic instability (Fig. S1A).

**Figure 1.**
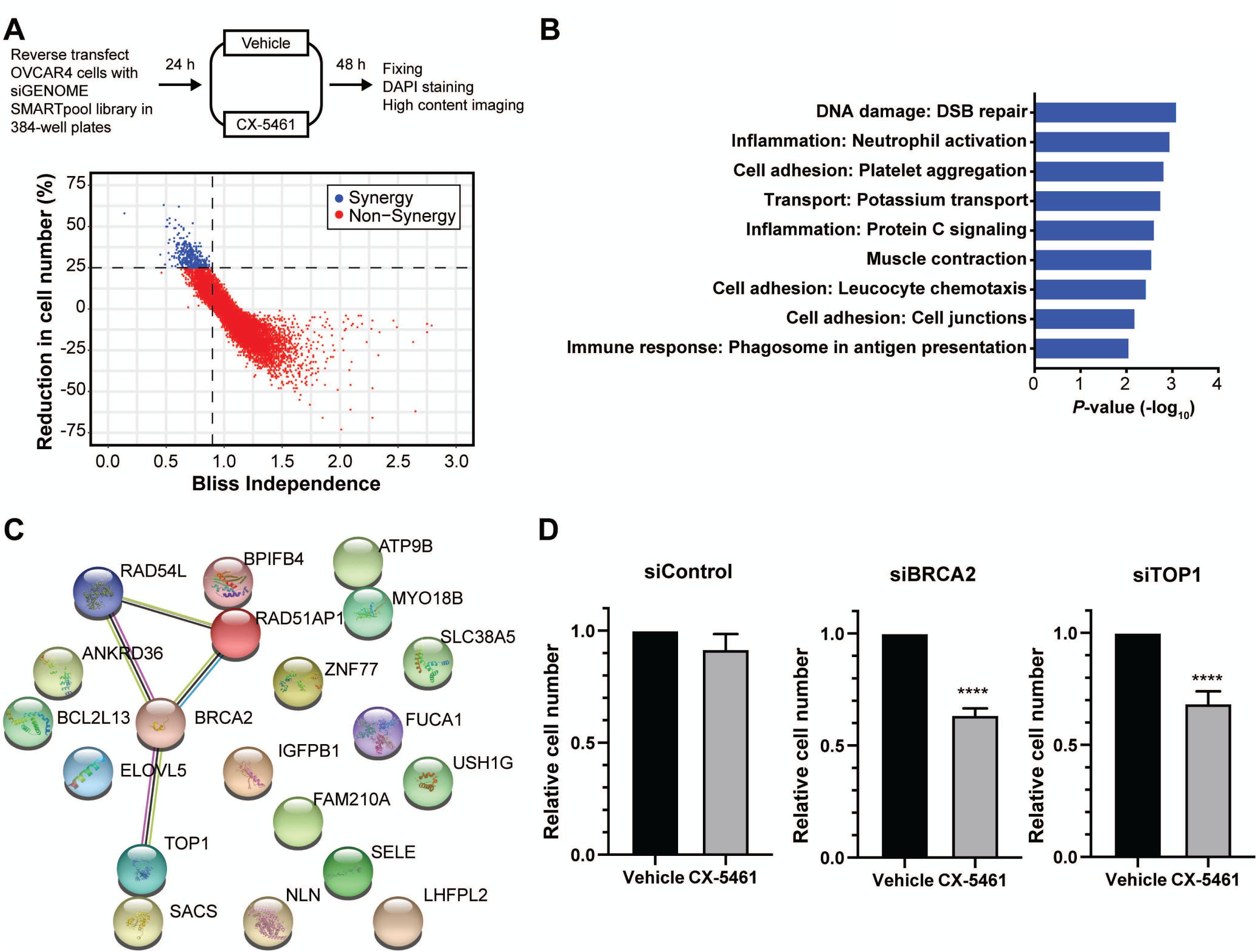
A genome-wide RNAi screen identifies a network of genes that when depleted synergise with CX-5461 in inhibiting HGSC cell proliferation. (**A)** Schematic representation of screen design. OVCAR4 cells were reverse-transfected with the siGENOME SMARTpool library. After 24 h cells were treated with vehicle control (400 nM NaH_2_PO_4_) or 80 nM CX-5461 (GI_25_ dose for proliferation at 48 h). At 48 h post-transfection, cells were fixed, DAPI stained and imaged by high content microscopy to determine cell number. (**B)** Significant enriched functional processes (*P*-value < 0.01) associated with the primary screen hits. (**C)** STRING network of the 20 validated screen hits. (**D)** Quantification of relative cell number for OVCAR4 cells transfected with control, BRCA2 or TOP1 siRNA and treated with vehicle or CX-5461 is presented as mean ± SEM and statistical significance was determined by two-tailed unpaired *t* test (****, *P* < 0.0001).

Of the validated candidates, we also identified DNA topoisomerase I (TOP1) (2/4 siRNA duplexes validated), whose physiological function is to relieve DNA supercoiling that occurs during transcription and replication by generating single-strand breaks (SSBs) ^21^. We confirmed TOP1 localization in the nucleus (Fig. S1B), and its depletion upon siRNA transfection led to a significant decrease in cell number with simultaneous CX-5461 treatment (Fig. 1D). TOP1 also colocalised with the upstream binding factor (UBF) to the nucleolar periphery (Fig. S1B) where rDNA is repaired, agreeing with our previous finding that CX-5461 induces nucleolar-specific defects in rDNA chromatin ^10, 12^.

### CX-5461 and topotecan synergise in inhibiting HGSC cell growth

Given that we observed combinatorial effects between CX-5461 and TOP1 depletion, and the TOP1 inhibitor topotecan is used in salvage therapy for recurrent HGSC, we hypothesised that CX-5461 would be able to extend topotecan’s utility. We investigated how topotecan cooperates with CX-5461 in a panel of three HR-proficient HGSC cell lines: OVCAR4, OVCAR3 and CAOV3. We observed drug synergy in all cell lines as quantified by Bliss Independence based on a reduction in cell number (Fig. 2A). Clonogenic assays of OVCAR4 cells demonstrated marked suppression of colony formation with the combination of CX-5461 and topotecan compared with either drug alone (Fig. S2A). Microscopic examination of these cells revealed an enlarged, flattened morphology consistent with a senescence-like phenotype. Indeed, real-time quantitative PCR (qPCR) demonstrated induction of the senescence-associated secretory phenotype (SASP) genes *IL1α*, *IL6* and *CXCL8* (Fig. S2B). We also observed p53-independent transcriptional induction of *CDNK1A*, which encodes for the cyclin-dependent kinase inhibitor p21. Furthermore, cell cycle analyses of OVCAR4, OVCAR3 and CAOV3 cells demonstrated significant arrest in the G2/M population (Fig. 2B,C). OVCAR4 and CAOV3 but not OVCAR3 cells also showed a significant increase in the Sub G1 dead cell population (Fig. 2D), which was partially rescued by the caspase inhibitor Q-VD-OPh, suggesting that the cell death induced by the combination, at least in part, is mediated through apoptosis. Together, these data demonstrate that the combination of CX-5461 and topotecan induces a senescence-like cell cycle arrest and cell death.

**Figure 2.**
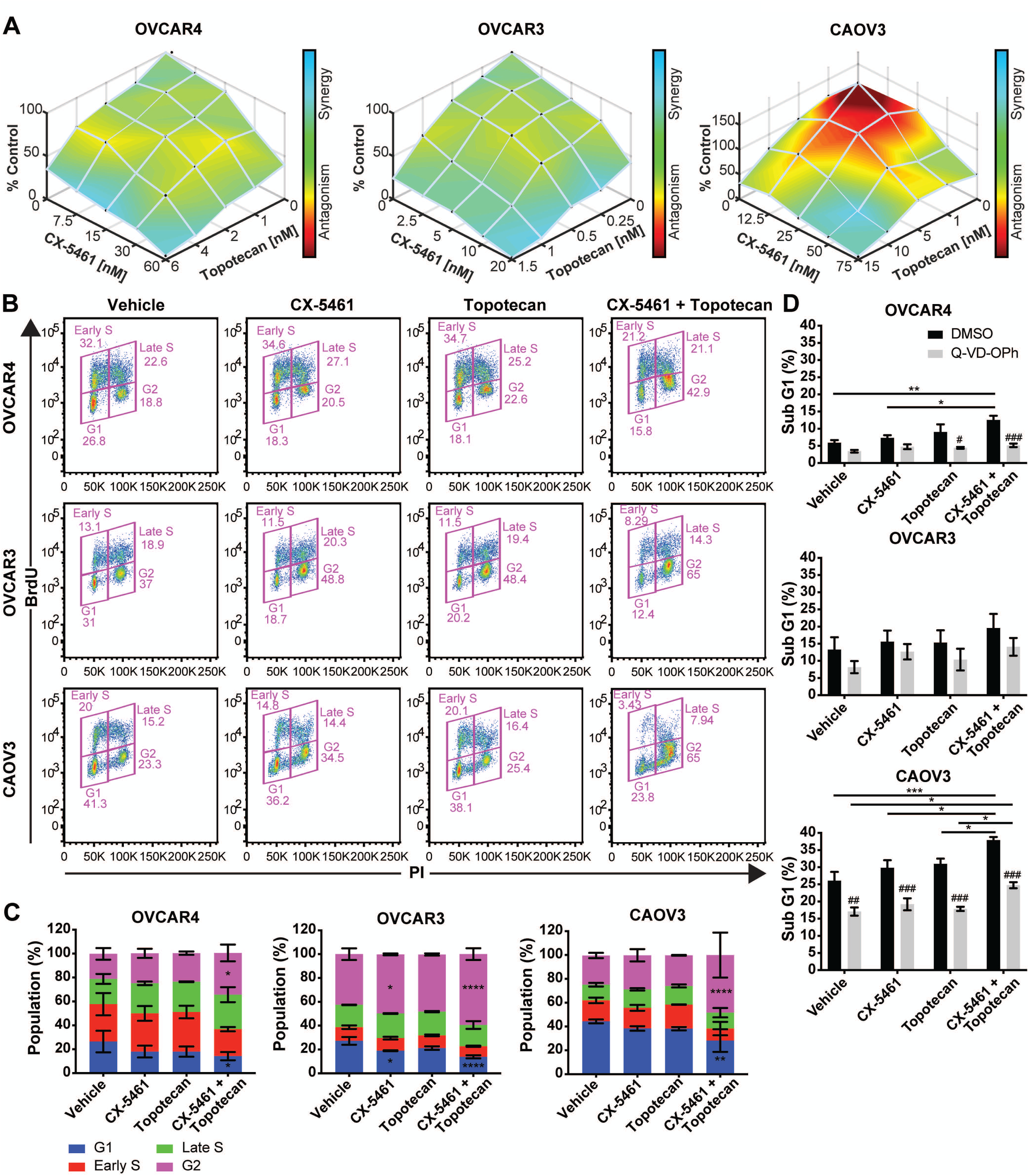
The TOP1 inhibitor topotecan synergises with CX-5461 in multiple HR-proficient HGSC cell lines. **(A)** 3D mapped surface BLISS plots for OVCAR4, OVCAR3 and CAOV3 cells treated with the indicated concentrations of CX-5461 and topotecan for 9 d. The range of concentrations for each drug was determined based on the GI_25_ at 5 d (Table S6). **(B)** FACS plots of cell cycle analysis of OVCAR4, OVCAR3 and CAOV3 cells treated with Vehicle, CX-5461 (80 nM, OVCAR4; 40 nM, OVCAR3; 360 nM, CAOV3; GI_25_ for proliferation at 48 h), Topotecan (6 nM, OVCAR4; 2 nM, OVCAR3; 15 nM, CAOV3; GI_25_ for proliferation at 48 h) or CX-5461 and Topotecan. **(C)** Quantification of the percentage of G1, Early S, Late S and G2 phase cells in **(B)** is presented as mean ± SEM and statistical significance was determined by Kruskal-Wallis one-way ANOVA (*, *P* < 0.05; **, *P* < 0.01; ****, *P* < 0.001; ****, P < 0.0001). (**D**) Quantification of the percentage of Sub G1 cells treated with control (DMSO) or 10 µM Q-VD-OPh is presented as mean ± SEM and statistical significance was determined by one-way ANOVA (*, *P* < 0.05; **, *P* < 0.01; ****, *P* < 0.001), (##, P < 0.01; ###, P < 0.001 for Q-VD-OPh as compared with DMSO control).

### The combination of CX-5461 and topotecan does not induce DNA double-strand breaks or further enhance replication stress

We previously showed the p53-independent response to CX-5461 is mediated by ATM and ATR, both of which are critical regulators of the DDR pathway ^10^ and are activated by various forms of DNA damage ^22^; hence, we hypothesised that the DDR would be a key mediator of the cellular response to CX-5461 plus topotecan. To test this hypothesis, we first assessed global DDR activation using Western blotting of phosphorylated checkpoint kinases (CHK)1 and CHK2 (downstream effectors of ATR and ATM, respectively), phospho-RPA (Ser33) (a marker of replication stress) and γH2AX (Ser139) (a marker of DNA damage) upon vehicle, CX-5461, topotecan or combination treatment for 3 or 24 h (Fig. 3A). We observed induction of phospho-CHK2 within 3 h and phospho-CHK1 by 24 h CX-5461 or topotecan treatment, which was markedly increased with the combination. At 24 h, in addition to phospho-CHK1 and phospho-CHK2, we also found a robust increase in phospho-RPA, indicating increased ssDNA. However, this did not translate to increased DSB formation, as we only observed γH2AX upon treatment with doxorubicin, a DNA topoisomerase 2 (TOP2) inhibitor known to cause DSBs, suggesting the combination of CX-5461 and topotecan may act via a distinct mechanism. Consistent with this hypothesis, alkaline comet assays of OVCAR4 cells revealed CX-5461 and topotecan as single agents or in combination did not induce comet tails, unlike doxorubicin (Fig. 3B). Thus, CX-5461, topotecan or the combination can induce the DDR without DNA resection.

**Figure 3.**
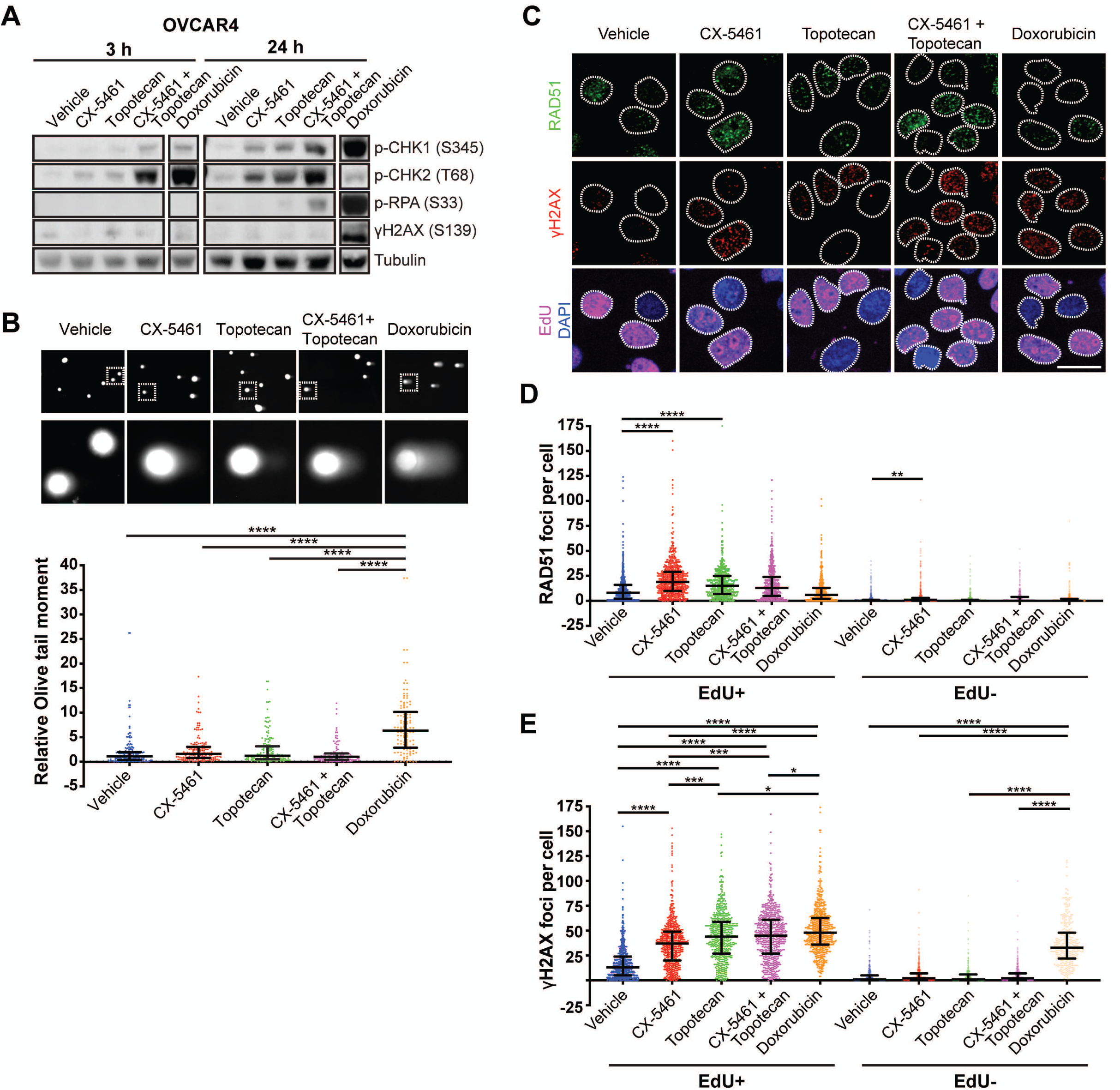
CX-5461 plus topotecan do not induce DNA resection or further enhance replication stress. **(A)** Representative Western blot analysis of DDR signalling in OVCAR4 cells treated with vehicle, 80 nM CX-5461, 6 nM topotecan, CX-5461 and topotecan or 1 μM doxorubicin for 3 h or 24 h (n=4 experiments). Tubulin was probed as a loading control. **(B)** Representative images and quantification of alkaline comet assay for OVCAR4 cells treated with vehicle, 1 μM CX-5461, 20 nM topotecan, CX-5461 and topotecan or 1 μM doxorubicin for 3 h is presented as median with interquartile range and statistical significance was determined by Kruskal-Wallis one-way ANOVA (**, *P* < 0.01; **, *P* < 0.0001). **(C-E)** OVCAR4 cells were treated with 1 μM CX-5461, 20 nM Topotecan, CX-5461 and Topotecan or 1 μM Doxorubicin for 3 h. **(C)** Representative images and (**D** and **E**) quantification of foci number for **(D)** RAD51 or **(E)** γH2AX. Cells were stained for EdU and DAPI to label replicating cells and nuclei, respectively. White dashed lines outline nuclei. Scale bar is 20 μm. Data are presented as median with interquartile range and statistical significance for increased foci per cell was determined by Kruskal-Wallis one-way ANOVA (*, P < 0.05, **, P < 0.01; ***, P < 0.001, ****, P < 0.0001).

We have shown that the CX-5461-mediated DDR is associated with degradation of stalled replication forks and replication-dependent DNA damage in HGSC cells ^12^. To determine if the combination of CX-5461 and topotecan induced further replication stress, we stained for RAD51 and γH2AX foci formation in OVCAR4 cells treated with 5-ethynyl-2’-deoxyuridine (EdU) to mark S-phase cells (Fig. 3C). Consistent with our previous finding, CX-5461 treatment increased RAD51 (Fig. 3D) and γH2AX (Fig. 3E) foci formation in EdU-positive cells compared to the EdU-negative cells, as expected for the induction of replication stress. Topotecan treatment also increased RAD51 and γH2AX foci formation; however, CX-5461 plus topotecan did not lead to a further enhancement.

### Combined treatment of CX-5461 and topotecan potentiates a nucleolar-associated DDR

As we have demonstrated CX-5461 alone induces the formation of ssDNA and recruitment of phosphorylated RPA to rDNA ^12^, we examined nucleolar recruitment of phosphorylated RPA (Fig. 4A) and found robust nucleolar localization of phospho-RPA with the combination treatment, particularly in replicating cells. We also examined the localization of phosphorylated ATR, which is recruited to RPA-coated ssDNA (Fig. 4B). Treatment with CX-5461 alone or in combination with topotecan enhanced nucleolar phospho-ATR staining as compared to vehicle. Furthermore, while nucleolar phospho-RPA was robustly induced upon doxorubicin treatment, phospho-ATR was not. Together, these data demonstrate that CX-5461 plus topotecan induces the RPA-ATR DDR axis within the nucleoli and a robust G2/M cell cycle arrest and cell death via a mechanism distinct from the standard DNA-damaging cytotoxic drug doxorubicin. Thus, we posit that a nucleolar-specific activation of the DDR will maintain treatment efficacy with reduced genotoxicity compared with standard cytotoxic drugs used to treat HGSC.

**Figure 4.**
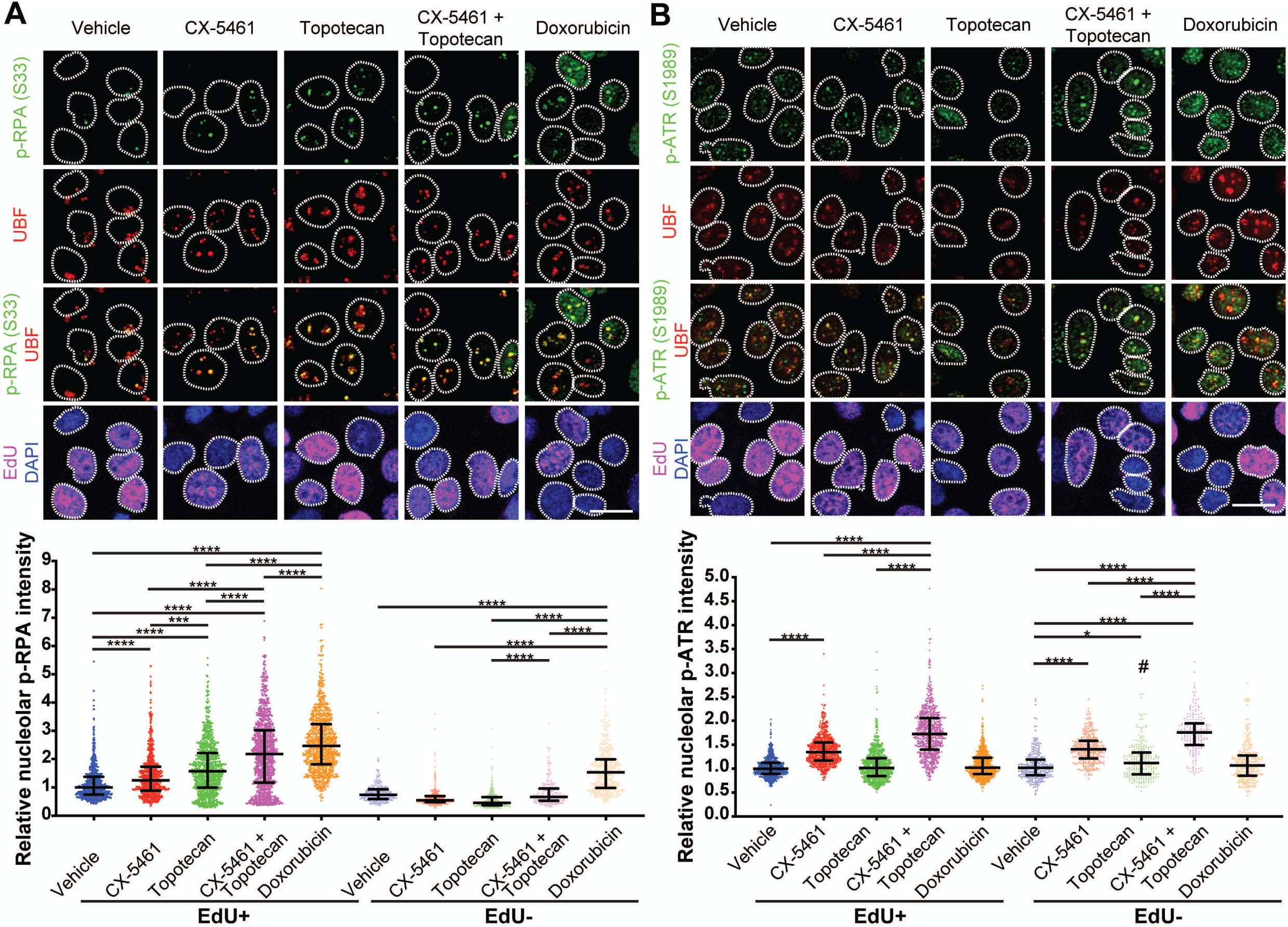
CX-5461 plus topotecan potentiates a nucleolar-associated DDR. (**A** and **B**) OVCAR4 cells treated with vehicle, 1 μM CX-5461, 20 nM topotecan, CX-5461 and topotecan or 1 μM doxorubicin for 3 h. Representative images and quantification of relative nucleolar fluorescence intensity for **(A)** p-RPA (S33) or **(B)** p-ATR (S1989). Cells were stained for UBF, DAPI or EdU to label nucleoli, nuclei or replicating cells, respectively. White dashed lines outline nuclei. Scale bar is 20 μm. Data are presented as median with interquartile range and statistical significance for increased p-RPA or p-ATR was determined by Kruskal-Wallis one-way ANOVA (*, *P* < 0.05; ***, *P* < 0.001; ****, *P* < 0.001), (#, P < 0.05; ####, P < 0.0001 for EdU-as compared with EdU+).

### CX-5461 plus topotecan delays HGSC xenograft progression

Topotecan is a primary salvage therapy for platinum-resistant HGSC but haematological suppression has restricted its long-term use in the clinic ^23^. Therefore, we hypothesised that the nucleolar-specific DDR induced by combining topotecan with CX-5461 would enable increased efficacy with less toxicity than single agent topotecan at the maximum tolerated dose (MTD). We established tumours from HR-proficient OVCAR3 cells harbouring amplified *CCNE1*, which confers primary treatment resistance and poor outcome of HGSC ^4^, and assessed tumour growth and survival in response to treatment with CX-5461 (30 mg/kg), topotecan (5 mg/kg) or CX-5461 plus topotecan (Fig. 5A). We chose a topotecan dosage of 5 mg/kg to mitigate the potential hematologic toxicity observed at higher doses (12.5 mg/kg) ^24^. Mice were dosed twice weekly for 4 weeks and then monitored until the animals reached an ethical endpoint. CX-5461 or topotecan treatment alone delayed tumour growth and prolonged survival as compared to vehicle treatment. Strikingly, the combined treatment of CX-5461 and topotecan blocked tumour progression during the entirety of drug treatment, extending survival. The combination was well tolerated with a mean weight loss < 10%. To test whether further reduction of the topotecan dose could retain therapeutic efficacy of the combination, we performed a second experiment using half the topotecan dose (2.5 mg/kg) of the first experiment (5 mg/kg) (Fig. 5B). In this setting, the combined treatment of CX-5461 and topotecan still significantly reduced tumour growth and extended overall survival compared to topotecan alone.

**Figure 5.**
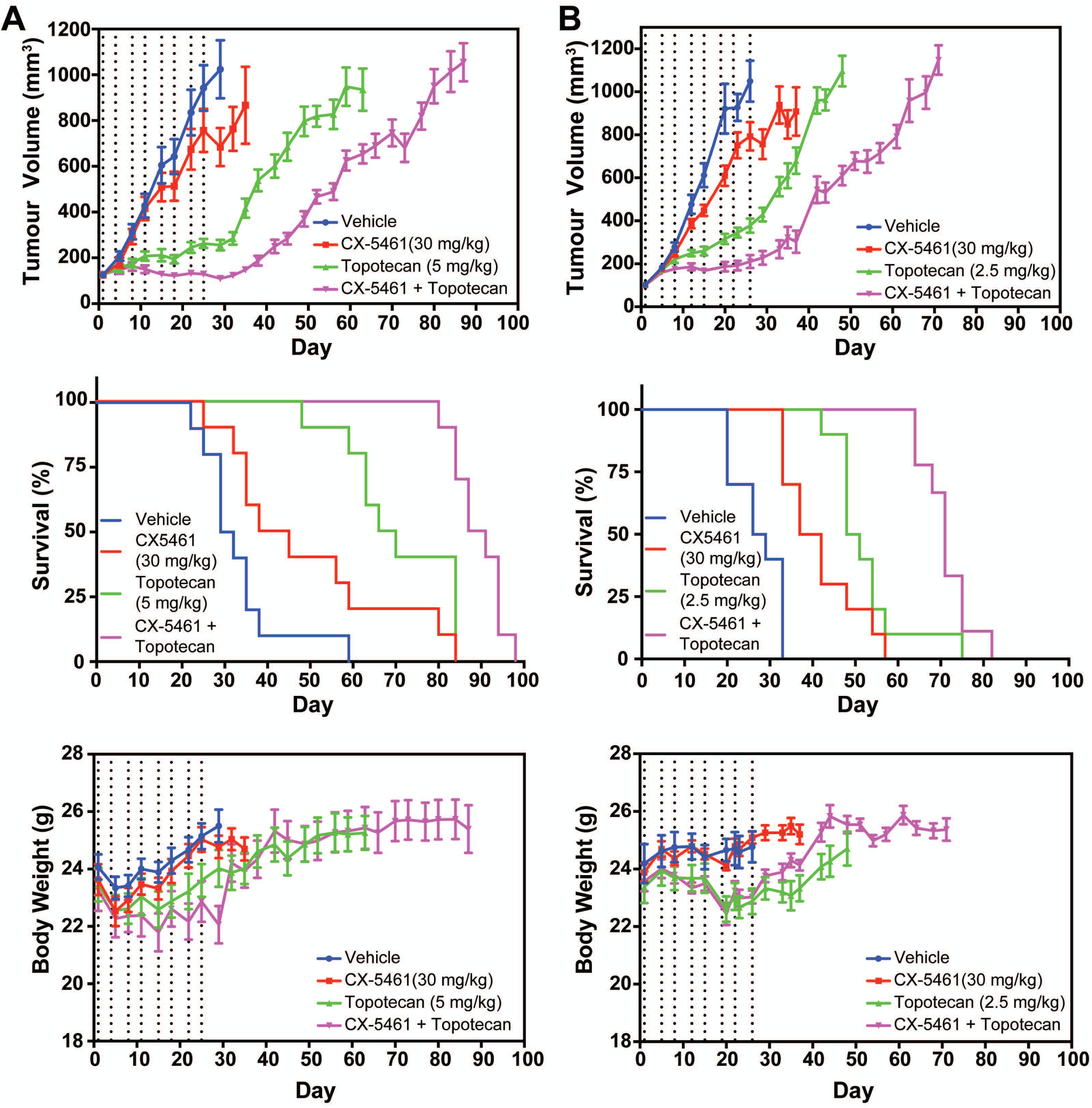
CX-5461 plus topotecan delays HGSC xenograft growth and prolongs survival. **(A)** (*top panel*) Mean tumour volume of OVCAR3 flank tumours treated with vehicle, CX-5461 (30 mg/kg), topotecan (5 mg/kg) or CX-5461 (30 mg/kg) and topotecan (5 mg/kg) (n=10 mice/group). Data are presented as mean ± SEM and statistical significance was determined on day 22 by one-way ANOVA (vehicle vs. CX-5461, topotecan or combination, CX-5461 vs. topotecan or combination, ****, *P* < 0.0001; topotecan vs. combination, ***, *P* < 0.001). (*middle panel*) Kaplan-Meier survival curves of tumour-bearing mice. Statistical significance was determined by log-rank Mantel-Cox tests (vehicle vs. CX-5461, CX-5461 vs. topotecan, *, *P* < 0.05; vehicle vs. topotecan or combination, CX-5461 vs. combination, ****, *P* < 0.0001; topotecan vs. combination, ***, *P* < 0.001). (*bottom panel*) Mouse body weight. **(B)** (*top panel*) Mean tumour volume of OVCAR3 flank tumours treated with vehicle (n=10 mice/group), CX-5461 (30 mg/kg) (n=10), topotecan (2.5 mg/kg) or CX-5461 (30 mg/kg) (n=10) and topotecan (2.5 mg/kg) (n=9). Data are presented as mean ± SEM and statistical significance was determined on day 20 by one-way ANOVA (vehicle vs. CX-5461, not significant; vehicle vs. topotecan or combination, CX-5461 vs. combination, ****, *P* < 0.0001; CX-5461 vs. topotecan, ***, *P* < 0.001; topotecan vs. combination, **, *P* < 0.01). (*middle panel*) Kaplan-Meier survival curves of tumour-bearing mice. Statistical significance was determined by log-rank Mantel-Cox tests (vehicle vs. CX-5461, *, *P* < 0.05; vehicle vs. topotecan or combination, CX-5461 vs. combination, ****, *P* < 0.0001; CX-5461 vs. topotecan, *, *P* < 0.05; topotecan vs. combination, ***, *P* < 0.001). (*bottom panel*) Mouse body weight. Dashed line indicates day of drug dosing.

## Discussion

We previously demonstrated that CX-5461 induces a nucleolar-associated DDR and is synthetic lethal with HR deficiency *in vitro* and *in vivo* in HGSC via a mechanism that is distinct from PARP inhibitors ^12^. Here, our genome-wide siRNA screen identified a number of genes involved in DNA repair (Table S4) including TOP1, which when depleted cooperate with CX-5461 to impair HGSC growth. We showed that CX-5461 the TOP1 inhibitor topotecan used in HGSC salvage therapy induces cell cycle arrest and cell death in multiple HGSC cell lines by potentiating a nucleolar-associated DDR, independent of DNA resection. The combination was also effective using low-dose topotecan in HGSC xenografts, thus providing a strategy to facilitate dose reduction to minimise the potential for toxicity while retaining efficacy to treat HGSC. In addition to confirming the importance of HR deficiency conferring sensitivity to CX-5461 in HGSC as we and others have shown ^12, 20^, we found TOP1 inhibition can cooperate with CX-5461 in the HR-proficient context, thereby potentially expanding the application of CX-5461 to a larger HGSC cohort, including patients previously harbouring HR-deficient tumours that have acquired chemotherapeutic resistance due to restored HR.

We demonstrated CX-5461 plus low-dose topotecan induces robust phosphorylation of RPA and ATR within the nucleolus. Together with our previous finding that CX-5461 enhances rDNA accessibility to MNase ^10^, we hypothesise that the combination further alters rDNA topology, which may occur via R loops ^12^ or G4-quadruplex DNA ^20^, to potentiate the DDR. Importantly, this nucleolar-associated DDR is less genotoxic than other DNA-damaging therapies such as doxorubicin, which does not activate nucleolar phospho-ATR but induces deleterious DSBs. Thus, CX-5461 plus topotecan represents a promising targeted approach for exploiting the nucleolar-associated DDR to improve salvage therapy.

Cellular senescence has been reported to be a central response to cytotoxic chemotherapy in HGSC ^25^. Indeed, we observed that a key cellular response of HGSC to CX-5461 plus topotecan is a G2/M cell cycle arrest and a senescence-like phenotype, which occurs through an enhanced nucleolar DDR without DNA resection. We posit that this can be further exploited in combination therapies with agents that selectively kill senescent cells ^26^ or engage anti-tumour immunity via the SASP, which can be further enhanced by agents promoting T cell activation to achieve even more durable responses. Overall, our findings highlight the potential of CX-5461 to enhance salvage therapeutic regimens for treating HGSC patients who have limited effective treatment options.

## Acknowledgments

We thank Dr. Daniel Thomas, Jennii Luu, Natalie Brajanovski, Jessica Ahern, Kerry Ardley and Rachael Walker for their technical support.

## Author Contributions

K.T.C. and R.B.P. conceived the project. K.T.C., R.B.P. and E.S. wrote the paper and S.Y., S.E., K.J.S., J.K., C.C., K.E.S., K.M.H. and R.D.H. contributed. S.Y. and K.T.C. performed the experiments. P.B.M. analysed the genome-wide RNAi screen data. Animal studies were performed in collaboration with C.C. The screen was a collaboration with K.J.S.

## Ethics declarations

### Competing interests

There are no competing financial interests in relation to the work described.

### Ethics approval

All animal experiments were performed in accordance with the Animal Experimentation Ethics Committee guidelines (Protocol #E557) at the Peter MacCallum Cancer Centre, Australia.

### Funding

The China Scholarship Council University of Melbourne PhD Scholarship supported S.Y. A National Health and Medical Research Council (NHMRC) Grant and NHMRC Senior Research Fellowship to R.B.P. supported this work. The Victorian Centre for Functional Genomics (K.J.S.) is funded by the Australian Cancer Research Foundation (ACRF), the Australian Phenomics Network (APN) through funding from the Australian Government’s National Collaborative Research Infrastructure Strategy (NCRIS) program, the Peter MacCallum Cancer Centre Foundation and the University of Melbourne Research Collaborative Infrastructure Program.

### Data Availability

The datasets are presented in the additional supporting files.

## References

1. Konstantinopoulos PA, Ceccaldi R, Shapiro GI, D’Andrea AD. Homologous Recombination Deficiency: Exploiting the Fundamental Vulnerability of Ovarian Cancer. Cancer Discov 2015; 5(11): 1137–1154; e-pub ahead of print 2015/10/16; doi 10.1158/2159-8290.CD-15-0714.

2. Ashworth A, Lord CJ. Synthetic lethal therapies for cancer: what’s next after PARP inhibitors? Nat Rev Clin Oncol 2018; 15(9): 564–576; e-pub ahead of print 2018/06/30; doi 10.1038/s41571-018-0055-6.

3. Christie EL, Fereday S, Doig K, Pattnaik S, Dawson SJ, Bowtell DDL. Reversion of BRCA1/2 Germline Mutations Detected in Circulating Tumor DNA From Patients With High-Grade Serous Ovarian Cancer. J Clin Oncol 2017; 35(12): 1274–1280; e-pub ahead of print 2017/04/18; doi 10.1200/JCO.2016.70.4627.

4. Au-Yeung G, Lang F, Azar WJ, Mitchell C, Jarman KE, Lackovic K et al. Selective Targeting of Cyclin E1-Amplified High-Grade Serous Ovarian Cancer by Cyclin-Dependent Kinase 2 and AKT Inhibition. Clin Cancer Res 2017; 23(7): 1862–1874; e-pub ahead of print 2016/09/25; doi 10.1158/1078-0432.CCR-16-0620.

5. Chekerov R, Hilpert F, Mahner S, El-Balat A, Harter P, De Gregorio N et al. Sorafenib plus topotecan versus placebo plus topotecan for platinum-resistant ovarian cancer (TRIAS): a multicentre, randomised, double-blind, placebo-controlled, phase 2 trial. Lancet Oncol 2018; 19(9): 1247–1258; e-pub ahead of print 2018/08/14; doi 10.1016/S1470-2045(18)30372-3.

6. Yan S, Frank D, Son J, Hannan KM, Hannan RD, Chan KT et al. The Potential of Targeting Ribosome Biogenesis in High-Grade Serous Ovarian Cancer. Int J Mol Sci 2017; 18(1); e-pub ahead of print 2017/01/25; doi 10.3390/ijms18010210.

7. Drygin D, Lin A, Bliesath J, Ho CB, O’Brien SE, Proffitt C et al. Targeting RNA polymerase I with an oral small molecule CX-5461 inhibits ribosomal RNA synthesis and solid tumor growth. Cancer Res 2011; 71(4): 1418–1430; e-pub ahead of print 2010/12/17; doi 10.1158/0008-5472.CAN-10-1728.

8. Khot A, Brajanovski N, Cameron DP, Hein N, Maclachlan KH, Sanij E et al. First-in-Human RNA Polymerase I Transcription Inhibitor CX-5461 in Patients with Advanced Hematologic Cancers: Results of a Phase I Dose-Escalation Study. Cancer Discov 2019; 9(8): 1036–1049; e-pub ahead of print 2019/05/17; doi 10.1158/2159-8290.CD-18-1455.

9. Hilton J, Cescon D, Bedard P, Ritter H, Tu D, Soong J et al. 44O CCTG IND. 231: A phase 1 trial evaluating CX-5461 in patients with advanced solid tumors. Annals of Oncology 2018; 29(suppl_3): mdy048. 003.

10. Quin J, Chan KT, Devlin JR, Cameron DP, Diesch J, Cullinane C et al. Inhibition of RNA polymerase I transcription initiation by CX-5461 activates non-canonical ATM/ATR signaling. Oncotarget 2016; 7(31): 49800–49818; e-pub ahead of print 2016/07/09; doi 10.18632/oncotarget.10452.

11. Gourley C, Balmaña J, Ledermann JA, Serra V, Dent R, Loibl S et al. Moving From Poly (ADP-Ribose) Polymerase Inhibition to Targeting DNA Repair and DNA Damage Response in Cancer Therapy. Journal of Clinical Oncology 2019: JCO. 18.02050.

12. Sanij E, Hannan KM, Yan S, Xuan X, Ahern JE, Chan KT, et al. Inhibition of RNA Polymerase I Transcription Activates Targeted DNA Damage Response and Enhances the Efficacy of PARP Inhibitors in High-Grade Serous Ovarian Cancer. bioRxiv 2019: 621623.

13. Domcke S, Sinha R, Levine DA, Sander C, Schultz N. Evaluating cell lines as tumour models by comparison of genomic profiles. Nat Commun 2013; 4: 2126; e-pub ahead of print 2013/07/11; doi 10.1038/ncomms3126.

14. Armstrong DK. Topotecan dosing guidelines in ovarian cancer: reduction and management of hematologic toxicity. The Oncologist 2004; 9(1): 33–42.

15. Bliss C. The toxicity of poisons applied jointly 1. Annals of applied biology 1939; 26(3): 585–615.

16. Schindelin J, Arganda-Carreras I, Frise E, Kaynig V, Longair M, Pietzsch T et al. Fiji: an open-source platform for biological-image analysis. Nature methods 2012; 9(7): 676.

17. Carpenter AE, Jones TR, Lamprecht MR, Clarke C, Kang IH, Friman O et al. CellProfiler: image analysis software for identifying and quantifying cell phenotypes. Genome biology 2006; 7(10): R100.

18. Gyori BM, Venkatachalam G, Thiagarajan P, Hsu D, Clement M-V. OpenComet: an automated tool for comet assay image analysis. Redox biology 2014; 2: 457–465.

19. Di Veroli GY, Fornari C, Wang D, Mollard S, Bramhall JL, Richards FM et al. Combenefit: an interactive platform for the analysis and visualization of drug combinations. Bioinformatics 2016; 32(18): 2866–2868.

20. Xu H, Di Antonio M, McKinney S, Mathew V, Ho B, O’Neil NJ et al. CX-5461 is a DNA G-quadruplex stabilizer with selective lethality in BRCA1/2 deficient tumours. Nat Commun 2017; **8**: 14432; e-pub ahead of print 2017/02/18; doi 10.1038/ncomms14432.

21. Pommier Y. Topoisomerase I inhibitors: camptothecins and beyond. Nature Reviews Cancer 2006; 6(10): 789.

22. Maréchal A, Zou L. DNA damage sensing by the ATM and ATR kinases. Cold Spring Harbor perspectives in biology 2013; 5(9): a012716.

23. McGuire WP, Blessing JA, Bookman MA, Lentz SS, Dunton CJ. Topotecan has substantial antitumor activity as first-line salvage therapy in platinum-sensitive epithelial ovarian carcinoma: a Gynecologic Oncology Group study. Journal of clinical oncology 2000; 18(5): 1062–1062.

24. Guichard S, Montazeri A, Chatelut E, Hennebelle I, Bugat R, Canal P. Schedule-dependent activity of topotecan in OVCAR-3 ovarian carcinoma xenograft: pharmacokinetic and pharmacodynamic evaluation. Clin Cancer Res 2001; 7(10): 3222–3228; e-pub ahead of print 2001/10/12.

25. Calvo L, Cheng S, Skulimowski M, Clement I, Portelance L, Zhan Y, et al. Cellular senescence is a central response to cytotoxic chemotherapy in high-grade serous ovarian cancer. bioRxiv 2018: 425199.

26. Fleury H, Malaquin N, Tu V, Gilbert S, Martinez A, Olivier M-A et al. Exploiting interconnected synthetic lethal interactions between PARP inhibition and cancer cell reversible senescence. Nature communications 2019; 10(1): 2556.

