## Supplementary Figure Legends for "Targeting RNA Polymerase I transcription synergises with TOP1 inhibition in potentiating the DNA damage response in high-grade serous ovarian cancer"

**Figure S1. Validation of BRCA2 or TOP1 knockdown.**

**(A)** OVCAR4 cells were transfected with control or BRCA2 siRNA for 48 h and treated with Vehicle or 80 nM CX-5461. After 48 h, cells were fixed and stained with DAPI to label cell nuclei. Quantification of percentage of cells with micronuclei is presented as mean ± SD and statistical significance was determined by Kruskal-Wallis one-way ANOVA (*, *P* < 0.05; ****, *P* < 0.0001). **(B)** OVCAR4 cells with transfected with control or TOP1 siRNA for 72 h and treated with Vehicle or 1 μM CX-5461 for 3 h. Cells were fixed and stained for TOP1, UBF as a nucleolar marker and DAPI to label nuclei. White dashed lines outline nuclei. Scale bar is 20 µm. Quantification of TOP1 intensity is presented as mean ± SD and statistical significance was determined by Kruskal-Wallis one-way ANOVA (****, *P* < 0.0001).

**Figure S2. CX-5461 plus Topotecan induce a senescence-like cell cycle arrest in HR-proficient HGSC.**

(**A)** Representative images of clonogenic survival assay for OVCAR4 cells treated with vehicle, 80 nM CX-5461, 6 nM topotecan or CX-5461 and topotecan for 48 h. Drugs were withdrawn and cells were cultured for an additional 5 d, fixed and stained with crystal violet. Quantification of percent confluence is presented as mean ± SEM and statistical significance was determined by one-way ANOVA (****, *P* < 0.0001). **(B)** Cells were treated as in **(A)** and qPCR was performed for the indicated genes. Quantification of relative RNA expression from is presented as mean ± SEM and statistical significance was determined by Kruskal-Wallis one-way ANOVA (*, *P* < 0.05; **, *P* < 0.01; ****, *P* < 0.001; ****, P < 0.0001 from n=3 experiments).
