## Supplementary figures and images for "Targeting RNA Polymerase I transcription synergises with TOP1 inhibition in potentiating the DNA damage response in high-grade serous ovarian cancer"

### Supplementary Figures S1 and S2

#### Yan et al. Figure S1

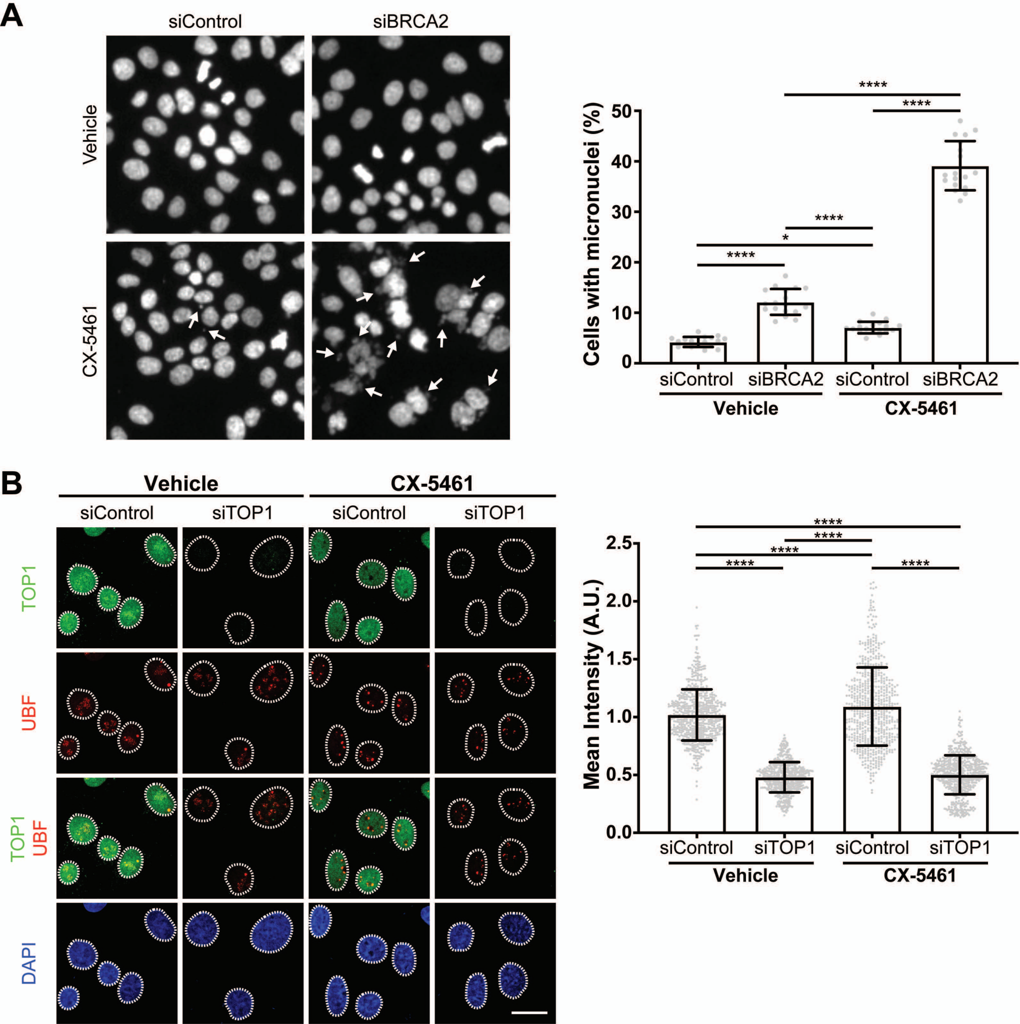

#### Yan et al. Figure S2

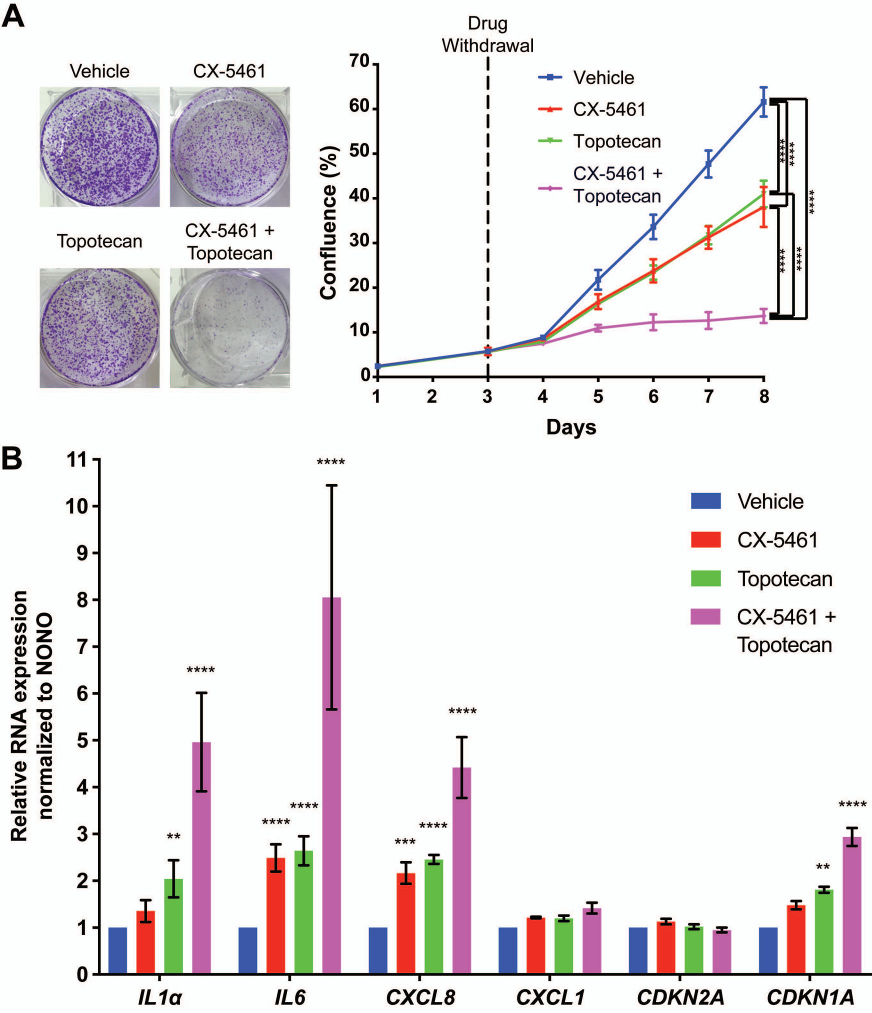
